# Meiotic drive, postzygotic isolation, and the Snowball Effect

**DOI:** 10.1101/2023.11.14.567107

**Authors:** Robert L. Unckless

**Affiliations:** Department of Molecular Biosciences, University of Kansas, Lawrence, KS 66045

**Keywords:** genetic conflict, toxin-antidote, gametic drive

## Abstract

As populations diverge, they accumulate incompatibilities which reduce gene flow and facilitate the formation of new species. Simple models suggest that the genes that cause Dobzhansky-Muller incompatibilities should accumulate at least as fast as the square of the number of substitutions between taxa, the so-called snowball effect. We show, however, that in the special— but possibly common— case in which hybrid sterility is due primarily to cryptic meiotic (gametic) drive, the number of genes that cause postzygotic isolation may increase nearly linearly with the number of substitutions between species.

## 1 Introduction

As populations diverge, they become increasingly reproductively isolated (reviewed in Coyne, Orr (2004)). The rate at which genes that cause hybrid incompatibilities accumulate between taxa has been the subject of much theoretical and empirical interest (Orr, 1995; Orr, Turelli, 2001; Kondrashov, 2003; Johnson, 2006; Moyle, Nakazato, 2008; Palmer, Feldman, 2009; Matute et al., 2010; Moyle, Nakazato, 2010; Wang et al., 2015; Dagilis et al., 2019; Kulmuni et al., 2020). Orr (1995) showed that under the traditional Dobzhansky-Muller (DM) model — in which any given substitution is potentially incompatible in hybrids with any previous substitution — the expected number of hybrid incompatibilities should increase at least as fast as the square of the number of substitutions between lineages. Orr referred to this accumulation as the “Snowball effect” and it occurs because each additional substitution is potentially incompatible with all prior substitutions. Several studies, however, have discussed scenarios in which incompatibility genes might instead increase linearly with the number of substitutions, e.g., Palmer, Feldman (2009); Kondrashov (2003); Dagilis et al. (2019). For present purposes, the most relevant of these scenarios is that in which a substitution is potentially incompatible only with a subset of previous substitutions which could lead to a more linear increase (Kondrashov, 2003). In this note, we highlight a biological scenario in which this condition may be satisfied: genetic conflict. Although genetic conflict has received a good deal of attention recently as a possible cause of hybrid problems (Presgraves, 2009; Phadnis, Orr, 2009; McDermott, Noor, 2010; Johnson, 2010; Crespi, Nosil, 2013; Presgraves, Meiklejohn, 2021), it has not, to our knowledge, been appreciated that conflict will not always yield a snowball effect.

The role of genetic conflict in speciation is perhaps unique as it can drag a population through a fitness valley— precisely the phenomenon that the Dobzhansky-Muller model was intended to avoid. Types of genetic conflict implicated in causing hybrid incompatibilities include meiotic/gametic drive (Tao et al., 2001; Phadnis, Orr, 2009; Presgraves, 2009; McDermott, Noor, 2010; Johnson, 2010; Lindholm et al., 2016), cytoplasmic male sterility (CMS) in plants (Fishman, Willis, 2006), and sexual conflict (reviewed in Arnqvist, Rowe (2005)). Here we focus on meiotic drive, first proposed by Frank (1991) and Hurst, Pomiankowski (1991). While these early models focused on the role that sex-ratio gametic drive (involving the sex chromosomes) might play in reproductive isolation, we focus on driving autosomes, for the sake of simplicity.

The role of genetic conflict in reproductive isolation is summarized in Johnson (2010) (see his Table 1) and Presgraves, Meiklejohn (2021). Of 18 characterized genes involved in reproductive incompatibilities, Johnson (2010) suggests that, in at least ten, conflict may have played a role in their divergence. And of six loci known to be involved specifically in hybrid male sterility, five may involve a history of conflict. Two of these genes (tmy and Ovd), both in Drosophila, are known to involve meiotic drive (Tao et al., 2007; Phadnis, Orr, 2009; Lai, Vogan, 2023).

Presgraves (2009) (p. 493) describes three ways in which gametic drive might lead to hybrid incompatibility:

1. The *strong model*, wherein hybrid sterility is caused by reciprocally-acting drive systems (after Frank (1991) and Hurst, Pomiankowski (1991)). Imagine a driving autosome that arises in population A and that goes to fixation. In population B, a *different* driving mutation arises on the same autosome and goes to fixation. Upon hybridization between the two populations, F1 males may well be sterile: if both drivers function in hybrids, the driver from population A will inactivate gametes bearing the autosomal homolog from population B and vice versa. The same result can occur if mutations that suppress gametic drive arise within each of the two populations and both the driving and suppressing mutations go to fixation (Hall, 2004). We refer to this scenario as mutually-assured destruction (MAD). Another flavor of the strong model is also possible. Imagine a single driver is unleashed in a hybrid. This would decrease the total number of gametes by about 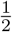, which may or may not cause a detectable reduction in male fertility. A second driver on a different chromosome (not the homolog in the other species) would reduce fertility further. The combined effects of two drivers would on average result in a ¾ reduction in gamete production. Three drivers on three different chromosomes would cause a 7/8 reduction and so on. We refer to this as the Incremental Reduction in Fertility (IRF) model. As soon as two drivers ended up on the same homolog in different species, however, the MAD model causes complete sterility.
2. The *weak model* (see also Frank (1991)), wherein divergence reflects an arms race between drivers and suppressors. In this case, a traditional Dobzhansky-Muller incompatibility can appear in hybrids as a result of this arms race. The non-independence of substitutions could be another reason for a linear increase in the number of incompatibility loci.
3. The *mixed model*, wherein a normally cryptic driver is released in hybrids and this allele both drives against the naïve genetic background of the other species (inactivating half of all gametes) and, suicidally, drives against itself (inactivating the other half). The result is hybrid sterility.

Because it is simple and has been the focus of most interest in the genetics of speciation, we consider *only* the strong model of meiotic drive. Our goal is merely to show that this scenario may lead to an increase in the number of loci causing hybrid incompatibilities that does not conform to the snowball pattern (increasing at least as fast as the square of substitutions).

## 2 Results

Under the standard DM model, the *K*th substitution is potentially incompatible in hybrids with all *K*-1 loci that diverged previously (Orr, 1995). The probability, *p*, that any two randomly chosen diverged loci are incompatible is assumed to be small. It follows, as Orr (1995) showed, that the expected number of traditional Dobzhansky-Muller incompatibilities (DMIs) is

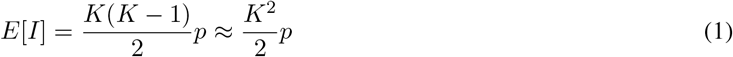

If substitutions obey a Poisson molecular clock, we can re-write Eq. (1) as a function of time. Orr, Turelli (2001) showed that

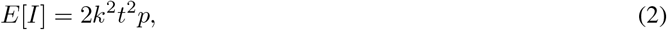

where *k* is the genomic rate of substitution of alleles, *t* is time, and this result is exact.

Under the meiotic drive scenario, an allele that drives is either suppressed in the same population in which it arose or sweeps to fixation without suppression – we do not allow stable polymorphism for drive. In either case, the driver eventually becomes fixed. We assume that if the driver is suppressed, the suppressor is recessive as drivers that are suppressed by dominant factors are unlikely to cause drive in hybrids. Upon hybridization, gametic drive is unmasked and gametes that are sensitive are killed.

The simplest model of meiotic drive causing incompatibilities assumes that drivers cause sterility with probability *p*_*d*_. In this case, the expected number of incompatibilities is simply *K*_*d*_*p*_*d*_, where *K*_*d*_ is the number of drivers that have arisen and either been fixed or suppressed. This simple model is obviously linear with *K*_*d*_.

In reality, a single driver may not cause discernible problems in hybrids. In many species, a 50% reduction in gametes count would likely have no noticeable effect on fertility. We explore two more complex models (MAD and IRF) that more realistically predict the effect of drive on fertility. In the mutually assured destruction model (MAD), different drivers fix on homologous chromosomes in two populations. Upon hybridization, males carry both drivers, leading to the destruction of all gametes.

### 2.1 Mutually Assured Destruction (MAD) Model

For the MAD model, there must be a driver on each homolog in the two diverged populations. The probability of this event depends on both the number of drivers that have fixed and the number of chromosomes in the species. We assume that both populations have the same chromosome complement and that all chromosomes are the same size. We relax the second assumption later.

To see the problem, imagine that a species has four chromosomes and that the first driver fixes in population A on chromosome 3. The probability that the second driver appears in population B on chromosome 3, causing mutually-assured destruction of gametes, is 1*/*8 = 1*/*(2*n*), where *n* is the number of chromosomes. From this it is clear that, for any number of drivers, MAD-related incompatibilities are more likely in species with fewer chromosomes.

Our interest is in the manner in which incompatibility loci accumulate over time, so we would like to determine the expected number of loci involved in incompatibilities after K drivers have fixed. We stress that we imagine an ideal experimental genetic analysis that can detect all loci involved in hybrid incompatibilities. This has two implications. First, it means that in those experiments, the phenotype assessed would be reduction in gametes as each driver alone only reduces gamete number by 1*/*2. Second, if a pair of homologous chromosomes contains three fixed drivers (two in one species and one in the other), all three are detected in our ideal genetic analysis and counted as incompatibility loci.

To derive the expected number of loci causing hybrid incompatibilities after *K* drivers fix, note that the number, *X*_*i*_, of drivers that fix on chromosome *i* = 1, 2, …, *n* is multinomially distributed, with 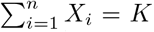. For clarity, consider only one of these *n* chromosomes. Some number, *X*, of the total *K* substitutions occur on this focal chromosome. *X* is binomially distributed: *X Binomial*(*K*, 1*/n*), *i*.*e*., with *K* trials and a probability of “success” of 1*/n*.

The *X* drivers that fix on the focal chromosome have two possible fates. Either they all land on the same homolog (all in population A or all in population B) or they are distributed over the homologs from the two populations (some in population A and some in population B). In the former case, *no* MAD incompatibilities arise on this chromosome and in the latter case *all X* substitutions act as MAD incompatibilities. The probability of the former event is 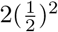 and, thus, the number of drivers that do not cause a MAD incompatibility on this chromosome is

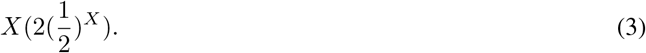

Averaging this quantity over the binomial distribution of *X*, the expected number of drivers that do not cause a MAD incompatibility on the focal chromosome is 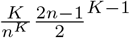.

To obtain the expected number of drivers on all chromosomes that are not involved in incompatibilities, we multiply by *n*, obtaining 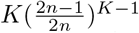. Consequently, the expected total number of MAD incompatibilities after *K* drivers is *K* minus the expected number that are not involved in MAD incompatibilities. This is

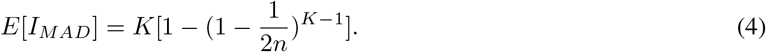

Figure 1 shows that, with small *n*, the increase in the number of incompatibility loci quickly approaches linearity. The reason is simple: once all chromosomes carry a MAD incompatibility, all subsequent drivers that fix on that chromosome (in either population) will also act as MAD incompatibilities (although no additional reduction in fertility occurs as fertility is already zero). The expected number of substitutions until saturation is 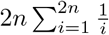, which for *n* = 2 is about eight drivers. With *n* = 10, however, it takes about 72 drivers to reach saturation.

**Figure 1.**
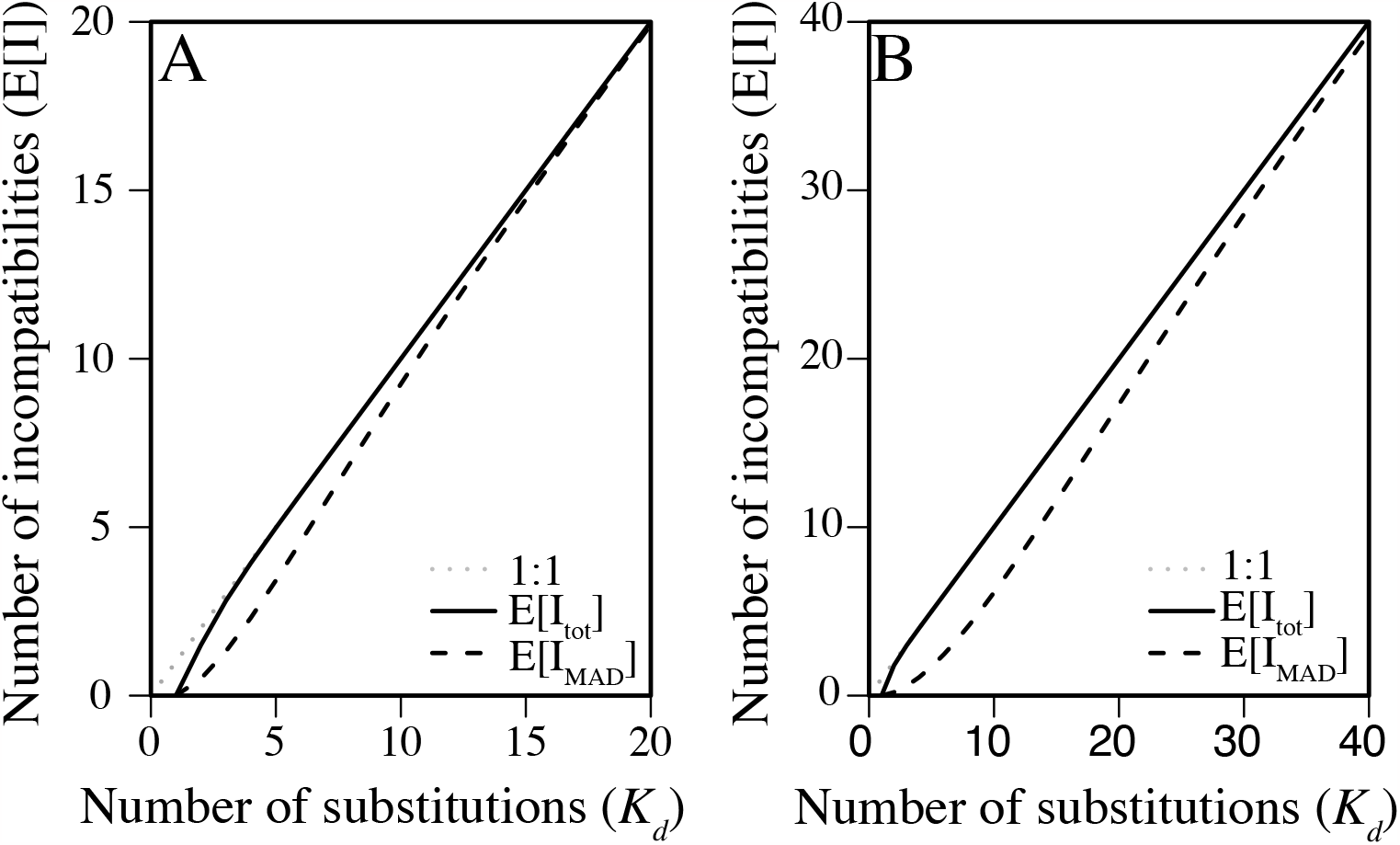
The accumulation of incompatibility caused by drive. Both the total number of incompatibilities and the number of MAD incompatibilities are presented as is a strict linear relationship. A) *n*=2 and B) *n*=10.

If chromosomes have different sizes, the result is only slightly more complex. Let *f*_*i*_ be the proportion of the genome represented by the *i*th chromosome such that ∑_*i*_ *f*_*i*_ = 1. Straightforward calculations show that the expected number of incompatibilities is now

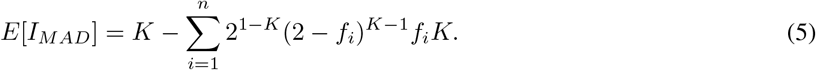

Using relative chromosome sizes from *D. melanogaster*, Eq. 5 leads to a linear increase in the number of MAD incompatibilities more quickly than under the assumption of equal chromosome size. In fact, taking the second derivative of Eq. 5 with n=2, shows that assuming equal chromosome sizes leads to the slowest approach to a linear increase.

While the MAD model eventually leads to an approximately linear increase in incompatibility loci, the number of drivers required to reach this point is large for even a moderate number of chromosomes.

### 2.2 Incremental Reduction in Fertility (IRF) Model

We now consider the incremental reduction in fertility model and the combined MAD and IRF models. Drivers causing an incremental reduction in fertility should increase nearly linearly at first, then disappear as chromosomes become saturated and the MAD model predominates. We must first arbitrarily decide at what amount of reduction in viable gametes is a fertility effect noticeable. It seems reasonable to assume that a 75% reduction in gametes could lead to noticeable effects, meaning that it would take two drivers on different chromosomes. While this number could be tuned, we use two drivers because it simplifies the analysis. Each additional driver causes some additional reduction in fertility as long as it does not land on a chromosome that already contains a driver (in either population).

So, after the first driver, each subsequent driver will cause an incremental reduction in fertility if it lands on any other chromosome in either species or mutually-assured destruction if it lands on the same chromosome in the other population. The only way it is not involved in sterility is if it lands on the *same* chromosome as a previous driver in the *same* population and not enough drivers have accumulated to produce a noticeable fertility effect.

To determine the expected number of total drive loci (MAD+IRF) involved in incompatibilities after *K*_*d*_ substitutions, we consider what happened with the previous *K*_*d*_−1 drivers. We are interested in two possible histories: *all* previous drivers land on the same chromosome in the same population or at least one landed on a different chromosome. In the combined model, the only way that a driver (after the first) does *not* cause an incompatibility is if it lands on the same chromosome in the same population as all previous drivers. A driver on the same chromosome as previous ones, but in the other population would cause a MAD incompatibility, while a driver on a different chromosome in either population would cause an IRF incompatibility. The probability that all *K*_*d*_ − 1 previous drivers land on the same chromosome is 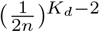. If the *K*_*d*_th driver lands on the same chromosome as all previous, we get no incompatibilities. If, however it lands on a different chromosome, there is an incompatibility and *all* previous drivers are now involved since, coupled with the *K*_*d*_th driver, any of the previous *K*_*d*_ − 1 could cause an incompatibility. The probability that the *K*_*d*_th driver lands on a different chromosome is 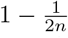 and if so, we must add *K*_*d*_ new incompatibility loci. The expected number of incompatibilities if the *K*_*d*_th driver is the first to cause an incompatibility is 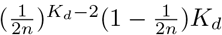.

If the previous *K*_*d*_ − 1 did not land on the same chromosome, then incompatibilities have already been established and the *K*_*d*_th driver adds one more. The expected number of incompatibilities at the *K*_*d*_th driver if at least one of the previous drivers landed on a different chromosome from the rest is 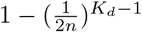. The total expected number of incompatibilities caused by the *K*_d_ th driver is the sum of the two expectations above: 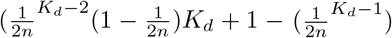. The expected total number of incompatibilities after *K* drivers is the sum of the previous over all drivers:

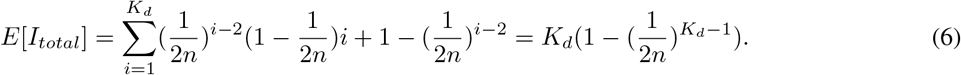

Figure 1 shows that the rate of increase of incompatibility loci caused by drivers approaches linearity by the second driver *regardless* of the number of chromosomes involved. It is also easy to see that the term in parentheses in equation 6 approaches one quickly. It is interesting to note, however, that the number of chromosomes in a species dictates which of the two models is likely to be more important in recently diverged pairs. If *n* is large, the probability of a MAD incompatibility is low since drivers will likely land on different chromosomes, therefore IRF incompatibilities play a larger role. Conversely, if *n* is small the probability of a MAD incompatibility is appreciable even after only two drivers have fixed.

If we relax the assumption that all chromosomes are the same size, as in equation 5, the total expected number of incompatibilities is now

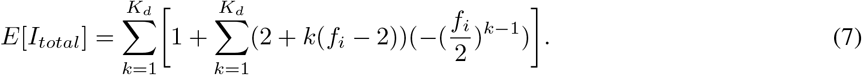

Interestingly, now equal sizes represents the fastest approach to linearity because the IRF model accumulates incompatibilities most quickly if all chromosomes are the same size.

### 2.3 Relative rates of DM incompatibilities compared to drive-related incompatibilities

Under most parameter values, a linear function increases faster than a power function until the point at which the power eclipses the linear increase. This is because while the slope of a straight line is constant, the slope of a power function increases over time. Models of Dobzhansky Muller and drive-related incompatibilities should show the same pattern. To determine how many substitutions occur before traditional DMIs overtake those caused by drive and therefore lead to the traditional snowball pattern, we must first define the proportion *p*_*d*_ of all substitutions that cause meiotic drive. This is analogous to *p* defined above that is the proportion of all substitutions that lead to incompatibilities. We can now write *K*_*d*_ as *K* ∗ *p*_*d*_ and solve for when Eqn 6 equals Eqn 1. This gives a simple result:

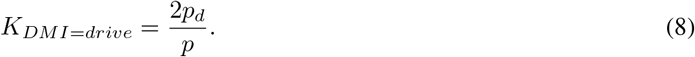

Even more simply, this can be stated as the twice the ratio of the likelihood that a substitution causes drive to the likelihood it causes a DM incompatibility. Figure 2 shows this simple relationship graphically. To observe the linear increase due to drive-related incompatibilities, we expect the equality in Eqn 8 to be at least 10 to 20, which would require that drive-related incompatibilities are ten times as likely as traditional DM incompatibilities.

**Figure 2.**
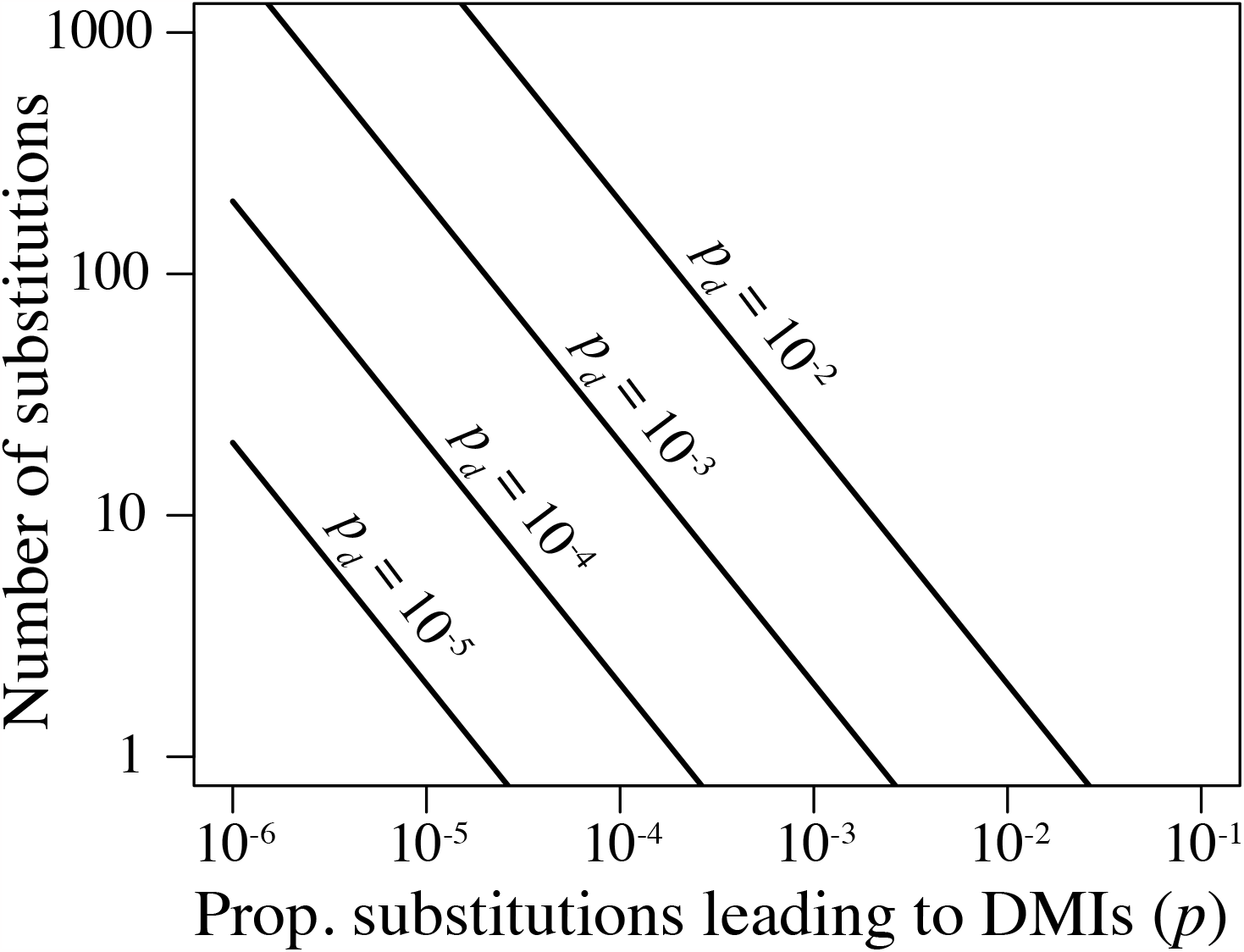
The number of substitutions where traditional DMIs overtake drive-related incompatibilities in total number. *p* is the proportion of all substitutions leading to DMIs and *p*_*d*_ is the proportion of all substitutions causing meiotic drive.

## 3 Discussion

Traditional models of the accumulation of hybrid incompatibilities assume that each substitution might be incompatible with all substitutions that occurred previously; this yields an approximately quadratic increase in the number of hybrid incompatibilities through time. Meiotic drive, on the other hand, may lead to a linear increase in hybrid incompatibilities. It is important to note, however, meiotic gametic drive is neither necessary nor sufficient for such a linear increase.

Several other models predict such linear increases; indeed our scenario represents a special, but possibly biologically important, case of a point made previously by Kondrashov (2003) and Gavrilets (2004). Gametic drive is also not sufficient to cause a linear increase as the weak and mixed models of Presgraves (2009) may yield a quadratic increase in the number of hybrid incompatibilities, as expected under the snowball process.

Unfortunately, few empirical studies have attempted to characterize the accumulation of hybrid incompatibilities in the correct manner— by counting the number of genes that cause incompatibilities between species pairs of different ages. (This is different, of course, from measuring the total strength of postzygotic isolation between species pairs.) A few empirical studies have, however, taken the required approach and provide mixed support for the snowball effect. Matute et al. (2010) find evidence of the snowball effect for hybrid inviability in Drosophila, whereas Moyle, Nakazato (2008) find evidence of the snowball effect for seed sterility in tomatoes. The latter authors could not, however, reject a linear accumulation of hybrid incompatibilities for pollen sterility in tomatoes. This latter result may be consistent with our results: if pollen sterility largely reflects a history of gametic drive, hybrid incompatibilities can increase linearly through time. (Needless to say, other explanations of this finding are possible.) Seed sterility is more complex as it may conflate true seed sterility with inviability of hybrid zygotes (L. Moyle, pers. comm.)— and hybrid inviability seems less plausible under the strong model of gametic drive. Finally, Wang et al. (2015) found evidence for the snowball on autosomes but not the X chromosome in house mice. They attribute this linear increase on the X to low mapping resolution, but the mutually assured destruction model could also explain the pattern.

When both traditional (DM) and drive-based hybrid incompatibilities occur, the chance of observing a snowball effect depends sensitively on the ratio of the fraction of substitutions that cause drive, *p*_*d*_, to the fraction of substitutions that cause DMIs, *p*. This fraction may, of course, vary among taxa and any estimate of its value is, at present, merely speculative. However, we initially discarded this hypothesis about the linear relationship between the accumulation of meiotic drivers and the snowball effect assuming that the proportion of substitutions necessary for observing a linear increase was simply too high to be realistic. It is hard to imagine the well-studied meiotic drivers like *segregation distorter* in *D. melanogaster* or the *t* haplotype in house mice arising *de novo* or moving around the genome fast enough to observe such a linear trend (Larracuente, Presgraves, 2012; Kelemen, Vicoso, 2018). These are large, co-adapted gene complexes with interacting parts and often a specific target on the homologous chromosome. The rampant discoveries of toxin-antidote meiotic drivers and their propensity for horizontal spread (Lai, Vogan, 2023; Eickbush et al., 2019; De Carvalho et al., 2022) suggest that such linear patterns in the accumulation of incompatibilities might be common. The *wtf* meiotic driver in fission yeast litters the genome, shows rapid evolution among paralogs, and, more often than not, the divergent copies do drive and are not able to suppress other paralogs (Eickbush et al., 2019). Copy numbers across *Schizosaccharomyces* are even more dynamic with an impressive 83 copies in *Schizosaccharomyces octosporus*. Likewise, the spore-killer meiotic driver, *Spok* litters the genome of *Podospera* and its relatives and likely moves horizontally through association with a transposon known as *Enterprise* (Vogan et al., 2021). *Drosophila simulans* hosts three different drive systems and one of them involves the gene *Dox* which, though not necessarily a toxin-antidote system, is mobile in the genome. Finally, genes involved in hybrid sterility between rice species *Oryza sativa* (Asian rice) and *O. glaberrima* (African rice), involve a toxin-antidote system (*S*1) (Xie et al., 2019). Thus, assuming that the fraction of mutations (*p*_*d*_) that cause meiotic drive is greater than the fraction that lead to traditional Dobzhansky Muller Incompatibilities *p* - the condition necessary for a linear increase in incompatibilities - may not be unreasonable.

Here we focused on meiotic drive as a cause of hybrid incompatibilities, but other types of genetic conflict may often yield a similar result. For example, a transposable element might spread by inserting itself throughout the genome but will ultimately be repressed by some mechanism (e.g., PIWI-piRNA, Aravin et al. (2007)). Upon hybridization, repression may fail and the element may insert freely— including into essential genes (see Kidwell et al. (1977); Erwin et al. (2015))— yielding hybrid incompatibilities. As the element is incompatible with only those loci involved in TE control, incompatibilities might increase linearly with time. Perhaps more directly related are the *Medea*-like toxin/antidote systems where toxin (and antidote) act zygotically, not in the gametes (Wade, Beeman, 1994). These systems appear to be quite common in multiple species of *Caenorhabditis* nematodes (Noble et al., 2021; Ben-David et al., 2021).

Our model is simple and a number of complications could be considered, including the roles of suppressors at loci other than the responder, of sex-chromosome distortion, of imperfect distorters, and of populations in which distorters attain an intermediate equilibrium frequency. While these complications may merit future investigation, we suspect that our qualitative findings will often (though surely not always) be unaffected. Imagine, for example, a recessive suppressor that fixes on a different chromosome from the driver/responder loci. Although the hybrid incompatibility now involves three loci, the driver will still inactivate gametes that carry a sensitive responder locus. Our algebra would not therefore change. So long as the strong model is considered and the original driver inactivates gametes that carry the sensitive responder, our model should hold, at least qualitatively.

## 4 Acknowledgements

H. Allen Orr was instrumental in putting together the original version of this manuscript. We thank S. Zanders, A. Gupta, J. Jaenike, D. Presgraves, M. Turelli, Y. Brandvain for very helpful comments and/or discussion. This work was supported by National Institutes of Health grant GM51932 to HAO and NSF CAREER 2047052 to RLU.

